# Expression interplay of calcium-binding genes and transcription factors during the osmotic phase provides insights on salt stress response mechanisms in bread wheat

**DOI:** 10.1101/2024.02.07.579402

**Authors:** Diana Duarte-Delgado, Inci Vogt, Said Dadshani, Jens Léon, Agim Ballvora

## Abstract

Bread wheat is an important crop for the human diet, but the increasing soil salinization is reducing the yield. The Ca^2+^ signaling events at the early stages of the osmotic phase of salt stress are crucial for the acclimation response of the plants through the performance of calcium-sensing proteins, which activate or repress transcription factors (TFs) that affect the expression of downstream genes. Physiological, genetic mapping, and transcriptomics studies performed with the contrasting genotypes Syn86 (synthetic, salt-susceptible) and Zentos (elite cultivar, salt-tolerant) were integrated to gain a comprehensive understanding of the salt stress response. The MACE (Massive Analysis of cDNA 3’-Ends) based transcriptome analysis until 4 h after stress exposure revealed among the salt-responsive genes, the over-representation of genes coding calcium-binding proteins. The functional and structural diversity within this category was studied and linked with the expression levels during the osmotic phase in the contrasting genotypes. The non-EF-hand category from calcium-binding genes was found to be specific for the susceptibility response. On the other side, the tolerant genotype was characterized by a faster and higher up-regulation of EF-hand genes, such as RBOHD orthologs, and TF members. This study suggests that the interplay of calcium-binding genes, WRKY, and AP2/ERF TF families in signaling pathways at the start of the osmotic phase can affect the expression of downstream genes. The identification of SNPs in promoter sequences and 3’-UTR regions provides insights into the molecular mechanisms controlling the differential expression of these genes through differential transcription factor binding affinity or altered mRNA stability.

**Key message:** The fine-tuned expression of calcium-binding genes and transcription factors during the osmotic phase underlies the susceptibility and tolerance to salt stress responses of contrasting bread wheat genotypes.

## Introduction

The increasing soil salinization worldwide is a major constraint for agriculture as most crops are salt-sensitive (Keshtehgar et al. 2013; Pessarakli and Szabolcs 2019). Bread wheat (*Triticum aestivum* L.) is a key staple crop for global food security that suffers significant reductions in yield and quality because of soil salinity (Zheng et al. 2009; Curtis and Halford 2014). The studies of complex traits in this species are challenging because of its intricate genome structure. This allohexapolyploid (2*n*=6*x*=42, AABBDD) contains three subgenomes of 16 Gb with 85% of repetitive sequences (Loginova and Silkova 2018; IWGSC 2018). Efforts from the International Wheat Genome Sequencing Consortium (IWGSC) have allowed the release of fully annotated and highly contiguous chromosome-level assemblies of diverse bread wheat lines (Montenegro et al. 2017; Shi and Ling 2018; IWGSC 2018; Borrill et al. 2019; Walkowiak et al. 2020; Zhu et al. 2021). The use of these assemblies has accelerated the dissection of complex traits as genes and genetic markers can be located in a physical context (Borrill *et al*., 2019). To achieve future global wheat demand, sequencing technologies can assist in the identification of molecular mechanisms, genes, and beneficial alleles for application in breeding programs aiming to develop cultivars with increased stress resilience (Varshney et al. 2021).

High soil salinity affects the plant growth because it leads to physiological drought conditions, ion toxicity, and cell oxidative damage (Tuteja 2007; Gupta and Huang 2014). According to the time of stress exposure, the plant growth response to salinity comprises the early osmotic and the late ionic phases. The osmotic phase is attributed to the reduced water potential in the rhizosphere because of the accumulation of salts in the soil and it is independent of the sodium accumulation in tissues (Ismail et al. 2014; Parihar et al. 2015; Julkowska and Testerink 2015). The primary consequence is the reduction of shoot growth and the production of new leaves because of the stomatal closure and the increase in leaf temperature (Roy et al. 2014). The signaling events in the osmotic phase include transient changes of intracellular Ca^2+^ that are sensed and provide crucial information for the acclimation response of the plants (Julkowska and Testerink, 2015). The elevation of [Ca^2+^]_cyt_ involves multiple channels, channel-like proteins, and pathways including calcium-binding genes regulating long-distance signaling and transcriptional reprogramming during stress (Himanen and Sistonen 2019; Malabarba et al. 2021). These calcium-binding genes have a diverse affinity for calcium ions. This affinity combined with their architecture and sub-cellular localization can contribute defining their biological roles in stress response pathways (Ranty et al. 2016). In addition, transcription factors (TFs) are regulating the expression of stress-responsive genes involved in calcium-related signaling pathways (Bürstenbinder et al. 2017; Wan et al. 2018). TFs can be activated or repressed in signal cascades triggered by Ca^2+^ and calcium sensors (Galon et al. 2010).

The functional and structural diversity within the calcium-binding category is broad and complex. Calcium-binding proteins are classified as EF-hand domain-containing or as non-EF-hand when other functional domains interact with Ca^2+^. Each one of these groups includes a variety of genes with diverse architectures and specific roles during susceptibility or tolerance responses to abiotic stresses (La Verde et al. 2018; Medvedev 2018; Mohanta et al. 2019). A better understanding of signaling pathways leading to abiotic stress tolerance in crop species can be obtained through the study of this gene category at the expression and sequence level. Natural variation in *cis*-regulatory sequences from calcium-binding genes can underlie the response to abiotic stresses by regulating mechanisms of expression in which TFs are involved (Ackermann et al. 2013; Cai et al. 2020).

To gain insights into the biological functions of genes and their influence on complex traits in wheat, the transcriptomic analyses during abiotic stresses (Sultana et al. 2020; Duarte-Delgado et al. 2020; Konstantinov et al. 2021) have been facilitated by the availability of a highly contiguous and fully annotated bread wheat genome assemblies (IWGSC 2018; Walkowiak et al. 2020). Previously, the Massive Analysis of cDNA 3’-ends (MACE) in leaves during the osmotic phase, revealed crucial differences in the expression profiles from the genes included in over-represented categories from contrasting bread wheat genotypes. For instance, the calcium-binding ontology was up- and down-regulated in the tolerant and susceptible genotypes, respectively (Duarte-Delgado et al., 2020).

The main goal of this research was to infer osmotic stress response mechanisms related to the expression of calcium-binding genes and the interplay with TFs in leaves from contrasting bread wheat genotypes. This study contributes to the understanding of the influence of calcium-related transcript expression and their related polymorphisms in the triggering of tolerance or susceptibility responses. This knowledge can be useful in the breeding of genotypes with improved resilience to salt stress effects. Some of the results are part of a PhD dissertation (Duarte-Delgado, 2020).

## Materials and Methods

### Clusters of time-course expression profiles

The expression of TFs and genes from the calcium-binding category during the osmotic phase was retrieved from the data collected by Duarte-Delgado et al. (2020) in two contrasting bread wheat genotypes. These genotypes consisted of the elite German winter wheat cultivar Zentos (salt-tolerant) and the synthetic genotype Syn86 (salt-susceptible), which are also contrasting parents from an AB-QTL study (Kunert et al. 2007; Dadshani, 2018). Clusters of time-course expression profiles were identified in the genes from the calcium-binding category, through the visualization of the patterns of up- or down-regulation at 8, 15, 30 min, and 4 h after stress exposure (ASE) in the two genotypes. The expression profiles from the TFs were clustered according to the family type and compared in the two genotypes. The LOESS (locally estimated scatterplot smoothing) option from the *ggplot2* library from R software (R Core Team 2021) was used to fit a curve representing the expression tendency of the members of each calcium-binding cluster and each TF family, as indicated by Duarte-Delgado et al. (2020).

### Classification of calcium-binding genes

The characterization of the functional domains from the salt-responsive genes included in the calcium-binding category was performed using the Interpro classification of protein sequences (Mitchell et al. 2019) available in the RefSeqv1.0 annotation of the wheat genome (Alaux et al. 2018). These genes were classified as EF-hand or non-EF-hand according to the calcium-binding domain type present.

### Phylogenetic analysis of EF-hand proteins and transcription factors

A comparative phylogenetic analysis of salt-responsive EF-hand proteins and TFs was performed with amino acid sequences from *Arabidopsis thaliana.* This analysis enabled the classification of the wheat EF-hand proteins without additional functional domains as calmodulin (CaM), calmodulin-like (CML), or calcineurin B-like (CBL) type. The EF-hand proteins with kinase domain were distinguished as members of the CPK (calcium-dependent protein kinases) family through the comparison with peptides sequences from CBL-interacting protein kinase (CIPK), CPK-related kinase (CRK), calcium/calmodulin-dependent protein kinase (CCaMK) and CPK members from *A. thaliana* (Ho 2015; Shi et al. 2018; Mohanta et al. 2019). Three NADPH oxidases (respiratory burst oxidase homologs, RBOHs) were also analyzed among the salt-responsive genes in wheat. These proteins with two EF-hand motifs, produce localized reactive oxygen species (ROS) bursts from the outer of the plasma membrane which are involved in the signaling to induce responses as stomatal closure upon stress sensing (Chapman et al. 2019; Shen et al. 2020). The genes from WRKY and APETALA2/Ethylene-Responsive-Element-Binding protein (AP2/ERF) families were further characterized and classified into subfamilies, as they were the most abundant TFs families in the transcriptomic analysis. A multiple sequence alignment was performed with MAFFT (Katoh et al. 2019) followed by a phylogenetic tree analysis conducted with MEGA X through the Neighbour-Joining method (Kumar et al. 2018). Consensus trees for salt-responsive CaM/CML, CPK, RBOH, AP2/ERF, and WRKY and reference proteins from *A. thaliana* were inferred through a bootstrap analysis with 3000 replicates. This phylogenetic analysis contributed to a better characterization of these salt-responsive genes, to deduce their functional properties and their putative involvement in regulatory networks related to stress response.

### RT-qPCR analysis of calcium-binding genes

The real-time quantitative PCR (RT-qPCR) analysis of two members of the RBOH family, *TraesCS5B02G299000* and *TraesCS4D02G324800*, was performed on leaves and roots in the time points studied in the transcriptomic analysis (Duarte-Delgado et al. 2020). Furthermore, the previous RT-qPCR data of the CML gene *TraesCS5D02G238700* on stressed leaves (Duarte-Delgado et al. 2020), was complemented with expression measurements in roots using the reported primers. Seeds of Zentos were purchased from Syngenta Seeds GmbH (Bad Salzuflen, Germany), while Syn86 seeds have been multiplied after their supply by Lange and Jochemsen (1992). Seedlings were in a growth chamber (20 ± 2 °C, 50 ± 5% humidity,12 h photoperiod) in the hydroponic system proposed and detailed by Dadshani (2018). Eight days after seed germination in Petri boxes with filter paper and distilled water, healthy seedlings were transferred to sponges located in panels over hydroponic boxes with 170 L of nutrient solution.

A salt treatment of 150 mM NaCl was applied to two-weeks-old seedlings that were sampled at 8 min, 15 min, 30 min, and 4 h ASE, according to the photosynthesis turning points previously identified (Dadshani 2018; Duarte Delgado et al. 2020). The control conditions corresponded to untreated hydroponics boxes containing plants harvested simultaneously with the stressed plants at the same time points. A biological replicate consisted of the whole root or all the leaves from a single plant. Three biological replicates were sampled from each organ, genotype, treatment, and time point. Leaves and roots were separated with scissors and immediately frozen in liquid nitrogen in 15 ml falcon tubes, followed by tissue homogenization with liquid nitrogen, mortar, and pestle. Total RNA isolation from 200 mg of grounded tissue was performed with the RNeasy plant mini kit (Qiagen, Hilden, Germany) and the cDNA synthesis with the First Strand cDNA Synthesis Kit (Thermo Scientific, Waltham, MA, USA) as described by Duarte-Delgado et al. (2020).

To design subgenome-specific primers for the salt-responsive RBOH genes (Table 1) the web-based tool GSP was used (Wang et al. 2016). Denaturation at 95 °C/7 min followed by 40 cycles at 95 °C/10 s, annealing temperature (Ta) for 30 s, 72 °C/30 s, and fluorescence acquisition temperature (Tfa) for 30 s were the cycling conditions used for RT-qPCR in a SDS-7500 Sequence Detection System device (Applied Biosystems, Waltham, MA, USA). Ta and Tfa conditions specific for each primer pair are listed in Table 1. The RT-qPCR reaction of 10 μl consisted of 0.25 μM of each primer, 5 µl of Luna^®^ Universal qPCR Master Mix (Biolabs, Ipswich, MA, USA) and 2 μl from 1:20 diluted cDNA template.

**Table 1.**
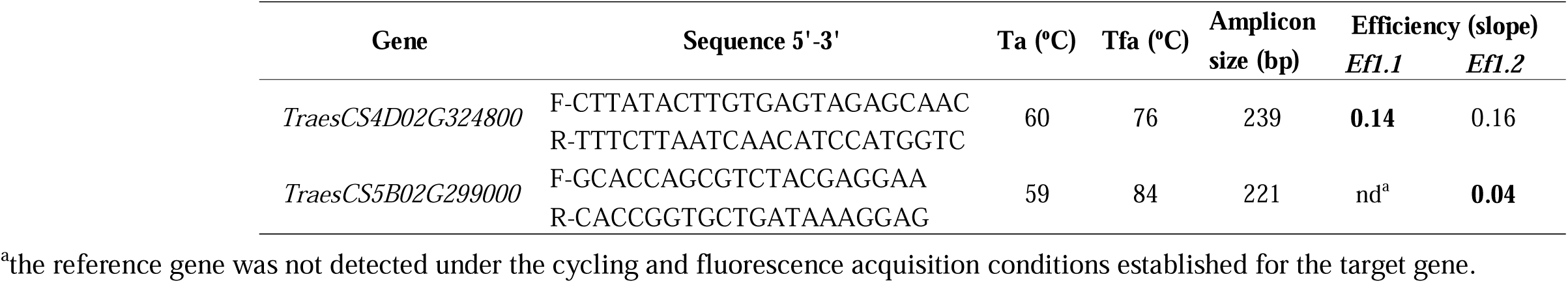
RT-qPCR primers of target genes with their respective annealing (Ta) and fluorescence acquisition (Tfa) temperatures. The amplification efficiency assessment was performed with the reference genes *Ef1.1* and *Ef1.2* (Oyiga et al. 2018, 2019) and was compared. Bold values represent the internal control selected for each target gene primer, according to the efficiency.

The amplification efficiencies of the internal control primers *TaEf-1.1* and *TaEf-1.2* (Oyiga et al. 2018, 2019) and each target gene were compared as proposed by Duarte-Delgado et al. (2020), and the control primer that resulted in the better efficiency was selected. For melting curve analysis (Supplementary Fig S1), a cycle of 95 °C for 10 s, Ta for 30 s, and 95 °C for 15 s was applied to PCR products. Based on the amplification efficiency comparison of the target and reference genes primers (Table 1), *TaEf-1.1* was the reference gene selected for the analysis of *TraesCS4D02G324800*. This reference gene was not detected in the RT-qPCR of *TraesCS5B02G299000*, because the Tfa defined in the cycling conditions (Table 1) was higher than the melting temperature from *TaEf-1.1* (82.5 °C; Duarte-Delgado et al. 2020). The other reference gene, *TaEf-1.2*, was therefore selected for the expression analysis in this case. The melting curves originated from the amplification of *TraesCS5B02G299000* revealed unspecific peaks (Supplementary Fig S1b). To avoid the contribution of the fluorescence from the unspecific products to the quantification, the temperatures of Tfa (Table 1) were adjusted to a temperature below the melting temperature (Tm) of the target product, as suggested by Klein et al. (2004).

The ΔΔ Ct method (Livak and Schmittgen 2001) was used to quantify the relative expression of the selected genes with the average Ct values of three technical replicates. A one-sample single-tailed t-test (*p*C<C0.05) was implemented in the 2^−ΔΔ*Ct*^ values, as detailed by Duarte-Delgado et al. (2020), to define whether the transcripts were up- or down-regulated upon stress in each time point, genotype, and organ. To determine if the mean relative expression values from both genotypes and organs were significantly different at each time point, a two-sample two-tailed t-test (*p*l*<*l0.05) was used to compare the 2^−ΔΔ*Ct*^ values from the same organ in the two genotypes and the two organs within the same genotype.

### Promoter sequence analyses

The promoter from *TraesCS2D02G173600, TraesCS5D02G238700,* and *TraesCS5B*02G299000 genes were analyzed by Sanger-based sequencing after PCR amplification. Subgenome-specific primers were designed (Table 2) as described in the previous section, based on the sequence of *ca.* 2000 bp upstream the start codon of the genes identified using the genome browser tool from the IWGSC RefSeq v1.0 (Alaux et al. 2018). The amplification conditions were adjusted using foliar DNA isolated with the Plant DNA mini kit (VWR, Darmstadt, Germany). Thus, 100 ng of DNA template in 25 µl of 1x One Taq Standard Buffer (Biolabs, Ipswich, MA, USA), 0.2 mM dNTPs, and 0.2 µM of each primer were amplified with 0.5 units of One Taq DNA polymerase (Biolabs, Ipswich, MA, USA). Cycling conditions were established with an initial denaturation step at 95°C/2 min followed by 40 cycles at 95 °C/45 s, Ta for 45 s (specified in Table 2), extension at 72 °C/1 min per kbp, and a final extension step at 72°C/5 min. The amplicons were visualized on 1.5% (w/v) agarose gels stained with peqGreen (0,04 µl/mL; VWR, Darmstadt, Germany). PCR products were purified with Purelink Quick PCR kit (Invitrogen, Waltham, MA, USA) and sequenced at Eurofins Genomics (Ebersberg, Germany) with an ABI 3730xl DNA Analyzer System.

**Table 2.**
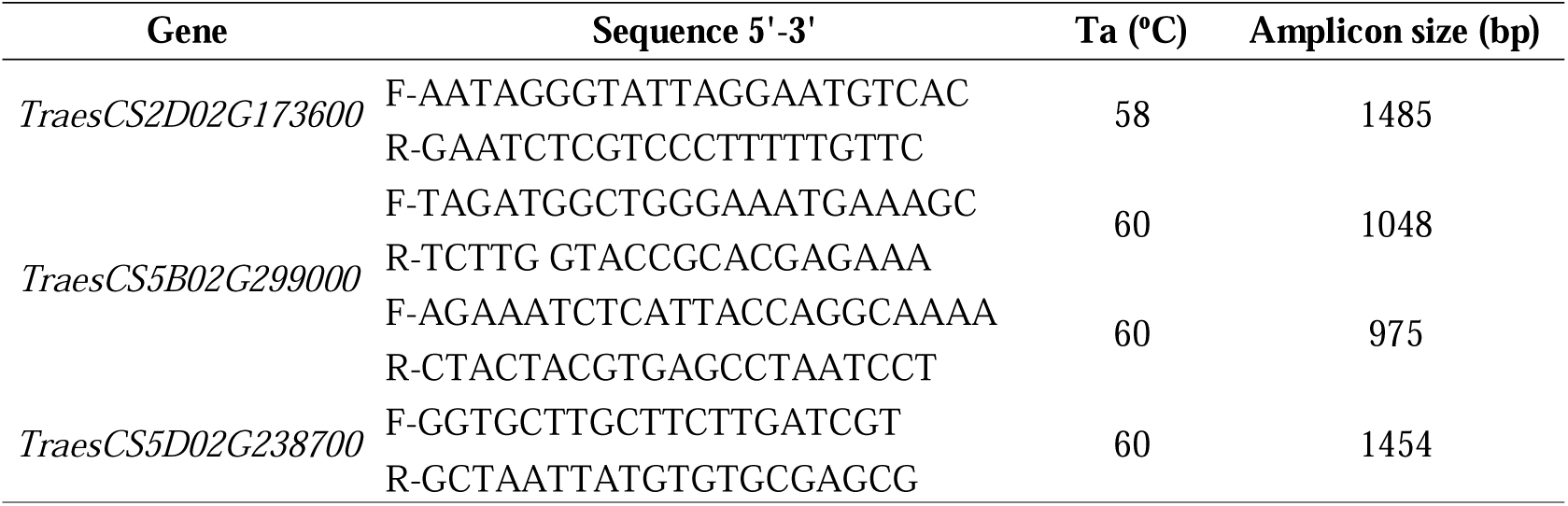
Primers designed to amplify the promotor regions of target genes studied by RT-qPCR, with their respective annealing temperatures (Ta).

Polymorphisms in the contrasting genotypes were identified using the pairwise alignment tool from Bioedit version 7.2.5 (Hall et al. 1999). The use of the PlantTFDB (v 5.0) database allowed the prediction of TF families with recognition motifs in the promoter sequences (Jin et al. 2017). TF families binding to motifs containing polymorphic sites were recognized as putative regulators of gene expression when *q-*value < 0.05.

### MACE-based SNP identification and miRNA binding site analysis

SNPs in salt-responsive calcium-binding genes were identified through the examination of the MACE reads collected for the transcriptomics analysis (Duarte-Delgado et al. 2020). These polymorphisms contributed to predict variations in miRNA binding sites that can be linked to the differences in expression observed in the contrasting genotypes (Võsa et al. 2015). The unique mapped reads from the MACE libraries of each genotype were merged for SNP identification using SAMtools (Li et al. 2009). Variant calling was performed with *samtools mpileup* followed by *bcftools call* commands from SAMtools, as described by Schneider et al. (2022). To retain high-confidence SNP calls, we established the following conditions: the minimum Phred base quality in the variant site was 25, the minimum mapping quality from the read was 20 and variants were represented in at least 10 reads from the same genotype. The psRNAtarget tool allowed the prediction of miRNA binding sites in the regions adjacent to the polymorphisms, based on known regulatory interactions in model species (Dai et al. 2018). Sequences of 10 nucleotides upstream and downstream the polymorphisms were retrieved for this analysis.

## Results

### Diversity of salt-responsive calcium-binding genes during the osmotic phase

The MACE analysis of leaves from 8 min to 4 h after salt stress exposure (ASE), detected different sets of salt-responsive genes within the calcium-binding category in Syn86 (salt-susceptible) and Zentos (salt-tolerant) (Duarte-Delgado et al. 2020). The proportion of salt-responsive genes coding EF-hand and non-EF-hand domains was determined in the contrasting genotypes (Fig. 1). The tolerant genotype showed 92% of calcium-binding genes containing EF-hand domain. From this, 76% were without any other functional domain and 12% contained a kinase domain. The non-EF-hand group was a minority (8%) with genes having an EGF-like domain (Fig. 1; Supplementary Table S1). A higher percentage of genes coding non-EF-hand domains was observed in Syn86 (53%). This genotype also showed increased functional and structural diversity within the EF-hand and non-EF-hand groups (Fig. 1; Supplementary Table S2). Most of the non-EF-hand type genes coded for oxygen-evolving complex (OEC) proteins (24%) followed by EGF-like domain proteins (13%). The EF-hand group consisted mainly of proteins that lacked any other functional domain (33%) followed by EF-hand proteins with other additional domains (5.5%). In the latter group, proteins such as caleosins (1.5%) and phosphoglycolate phosphatases (2%) were included (Supplementary Table S2).

**Fig. 1.**
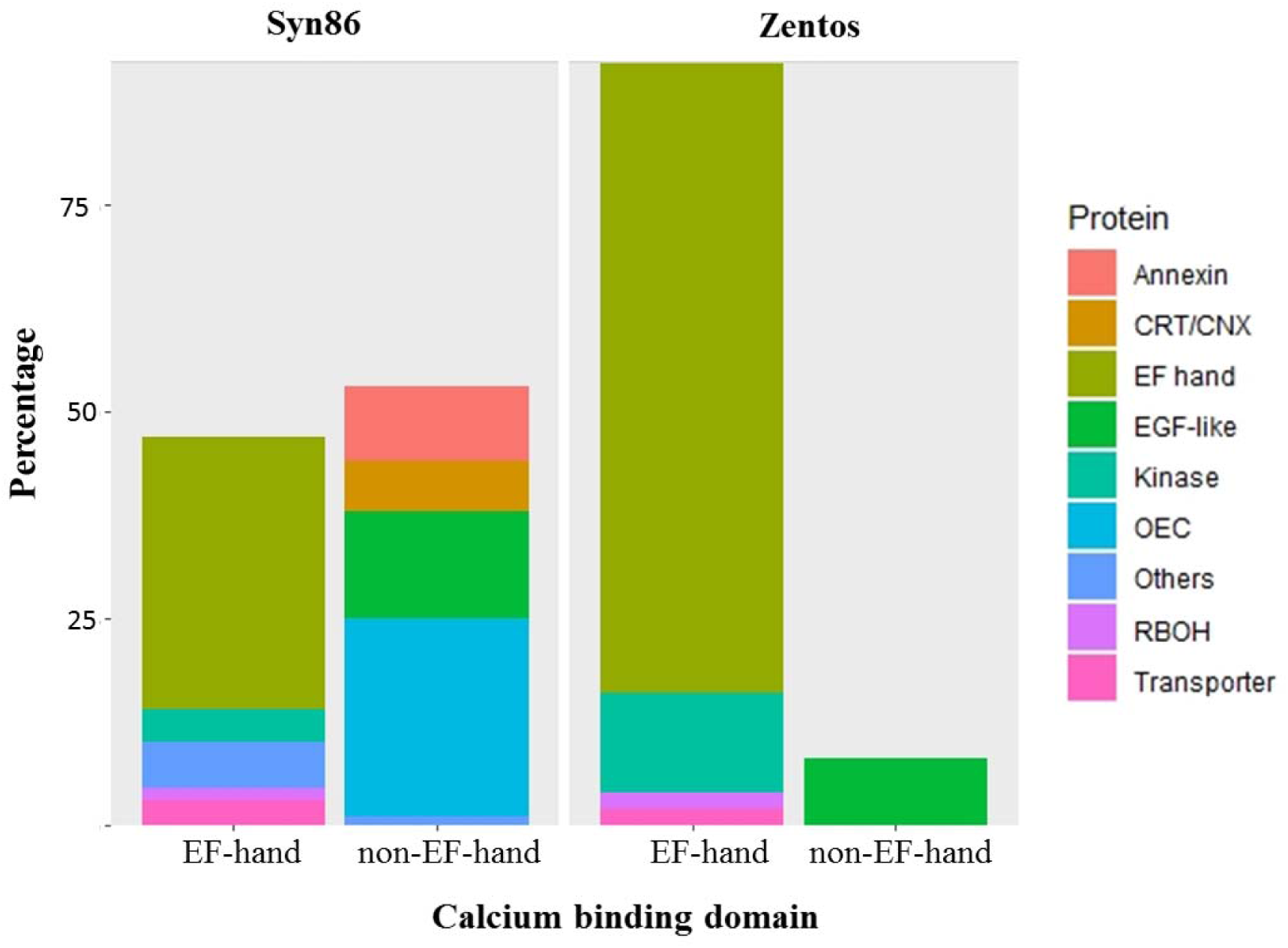
Distribution of EF-hand and non-EF-hand proteins coded by the salt-responsive calcium-binding genes identified in the contrasting genotypes.

The phylogenetic analysis of EF-hand domain proteins revealed the similarities and the clustering with CaM/CML, CBL, CPK, and RBOH members from *A. thaliana.* The subgroup of CaM genes had the highest statistical support (98%) in the CaM/CML analysis and included five salt-responsive genes from Syn86 (Fig. 2). From all proteins containing an EF-hand domain, 57 (72%) corresponded to the CML type. Mostly low bootstrap values (<50%) were observed in the corresponding clusters (Fig. 2). This analysis therefore revealed orthologous relationships with low statistical support among wheat and *Arabidopsis* CML sequences. The salt-responsive *TraesCS1B02G370900* from Syn86 was the only transcript identified from the CBL type and is orthologous to *CBL8*.

**Fig. 2.**
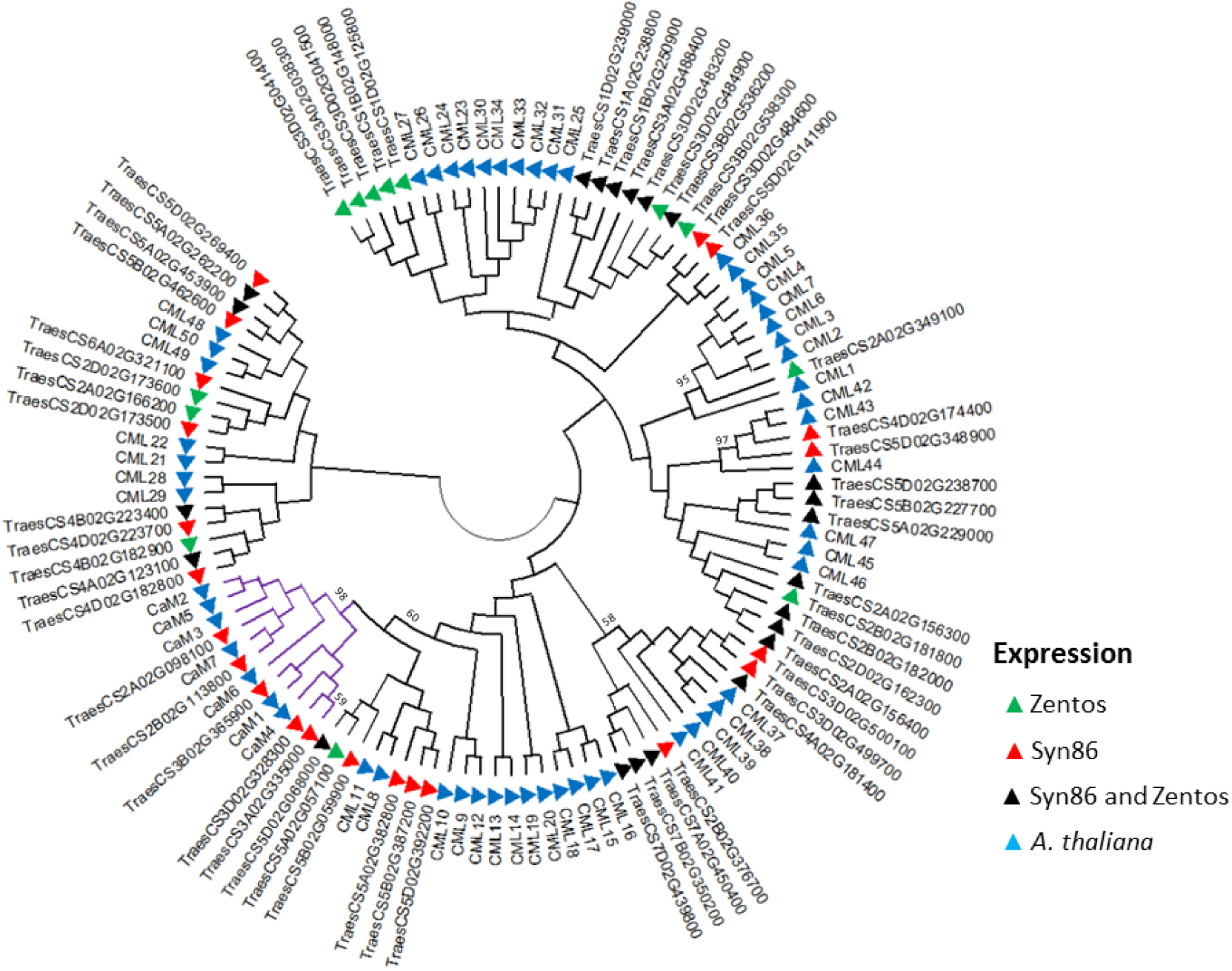
Dendrogram of CaMs (purple) and CMLs (black) amino acid sequences from *Arabidopsis thaliana* and coded by the corresponding salt-responsive genes from Zentos and Syn86 indicated by the colored triangles. The consensus phylogenetic tree was constructed with MEGA X (Kumar et al., 2018) using the Neighbour-Joining method and through a bootstrap analysis of 3000 replicates. Clusters of wheat and *Arabidopsis* proteins with bootstrap values > 50 are shown.

All the differentially expressed genes coding EF-hand proteins with kinase domain were from the CPK type, and clustered with good statistical support in subgroups I, III, and IV based on the classification proposed by Yip-Delormel and Boudsocq (2019) (Fig. 3). Three wheat genes were found in the subgroup I. From them, two homoeologous genes located in chromosomes 5A and 5B were up-regulated in Zentos at 15 min and were included in a cluster with *CPK1*, *CPK2,* and *CPK20*. The subgroups III and IV contained two and three wheat genes, respectively (Fig. 3). Similarly, the analysis of RBOH proteins revealed clusters with good statistical support (Fig. 4). From three proteins of wheat, two clustered with RBOHD and the third presented similarity with RBOHB. The wheat RBOHD orthologs were up-regulated, *TraesCS4D02G324800* in Zentos at 15 min and *TraesCS5B02G299000* in Syn86 at 30 min.

**Fig. 3.**
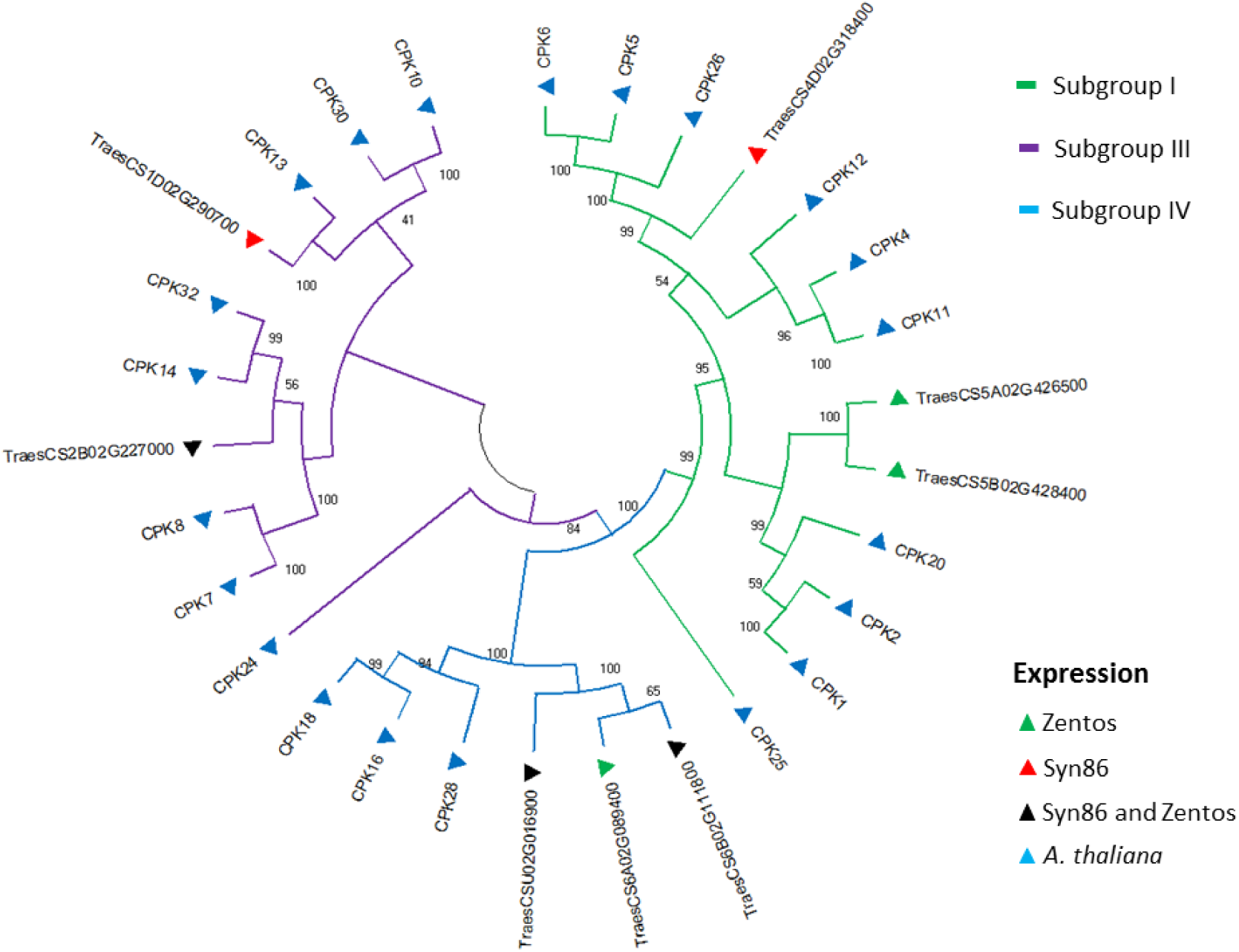
Dendrogram of CPKs amino acid sequences from *Arabidopsis thaliana* and coded by the corresponding salt-responsive genes from Zentos and Syn86 indicated by the colored triangles. The consensus phylogenetic tree was constructed with MEGA X (Kumar et al. 2018) using the Neighbour-Joining method and through a bootstrap analysis of 3000 replicates. The subgroups indicated by colors in the branches were defined according to the classification suggested by Yip-Delormel and Boudsocq (2019).

**Fig. 4.**
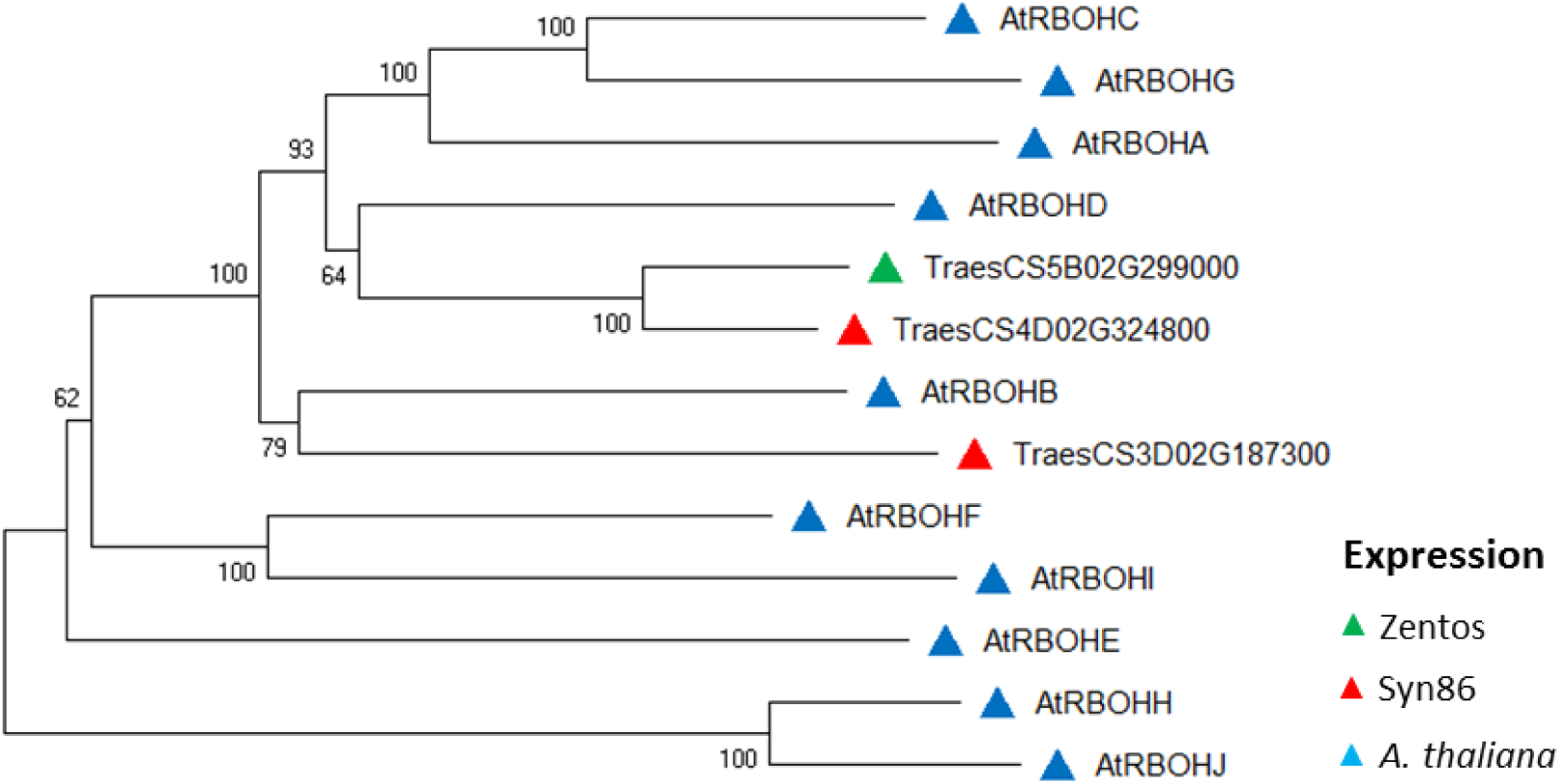
Dendrogram of RBOHs amino acid sequences from *Arabidopsis thaliana* and the corresponding salt-responsive genes from Zentos and Syn86 indicated by the colored triangles. The consensus phylogenetic tree was constructed with MEGA X (Kumar et al. 2018) using the Neighbour-Joining method and through a bootstrap analysis of 3000 replicates.

Clusters of time-course expression profiles were determined for the salt-responsive calcium-binding genes from each genotype (Fig. 5). The tolerant genotype showed five clusters, the main one with genes up-regulated at 15 min followed by a group of transcripts up-regulated both at 15 and 30 min (Fig. 5A). Seven clusters were defined in the susceptible genotype. Clusters I and II grouped the highest number of genes with down- and up-regulated transcripts at 30 min, respectively (Fig. 5B). The cluster I contained mainly OEC proteins and annexins, while the cluster II included EF-hand domain, EGF-like and calnexin/calreticulin (CNX/CRT) proteins. The two genotypes shared 27 calcium-binding genes, 22 of which corresponded to the CML type. The up-regulated transcripts across the stress response showed greater relative expression values in Zentos compared to Syn86 (Fig. 5C). At 4 h ASE, 13 of these genes were down-regulated in the susceptible genotype. The time-course expression values, the annotations, and the clusters of each calcium-binding gene are included in Supplementary Tables S1 and S2.

**Fig. 5.**
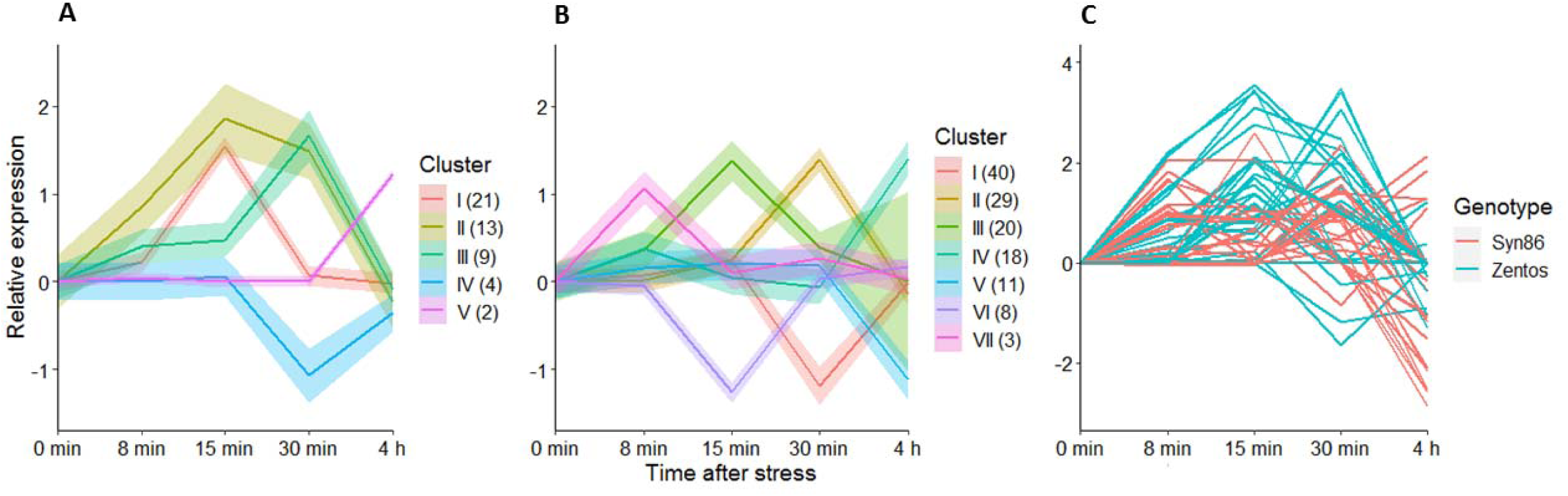
Expression profiles of salt-responsive genes with calcium-binding domain at the osmotic phase grouped in clusters represented by Roman numbers. The number of genes in each cluster is in parenthesis. Clusters from Zentos (A) and Syn86 (B) in which a curve was fitted to represent the expression tendency of the transcripts from each group and the shadows indicate the standard error of the relative expression values. Expression profiles of 27 calcium-binding transcripts identified in both genotypes (C).

### Time-course expression of EF-hand genes in roots and leaves by RT-qPCR

The time-course expression profiles of one CML member (*TraesCS5D02G238700*) and two members from the RBOH family (T*raesCS5B02G299000* and *TraesCS4D02G324800*) were analyzed by RT-qPCR in leaves and roots. The time-course analysis of the RBOH family members indicated that transcripts were up-regulated earlier in leaves than in roots. The measurements of *TraesCS4D02G324800* in leaves revealed the transcript up-regulation in Zentos at 8 min and in both genotypes at 15 min ASE. At this later time point, the relative expression values were the highest and there were non-significant differences in the mean values from the two genotypes. At 30 min, the gene was only up-regulated in Syn86, while at 4h it was downregulated in both genotypes (Fig. 6A). The analysis in roots showed the transcript up-regulation in Syn86 at 15 min and in both genotypes at 30 min ASE without significant differences (Fig. 6A). The mean expression values from the two organs were different in the tolerant genotype across all the time points.

**Fig. 6.**
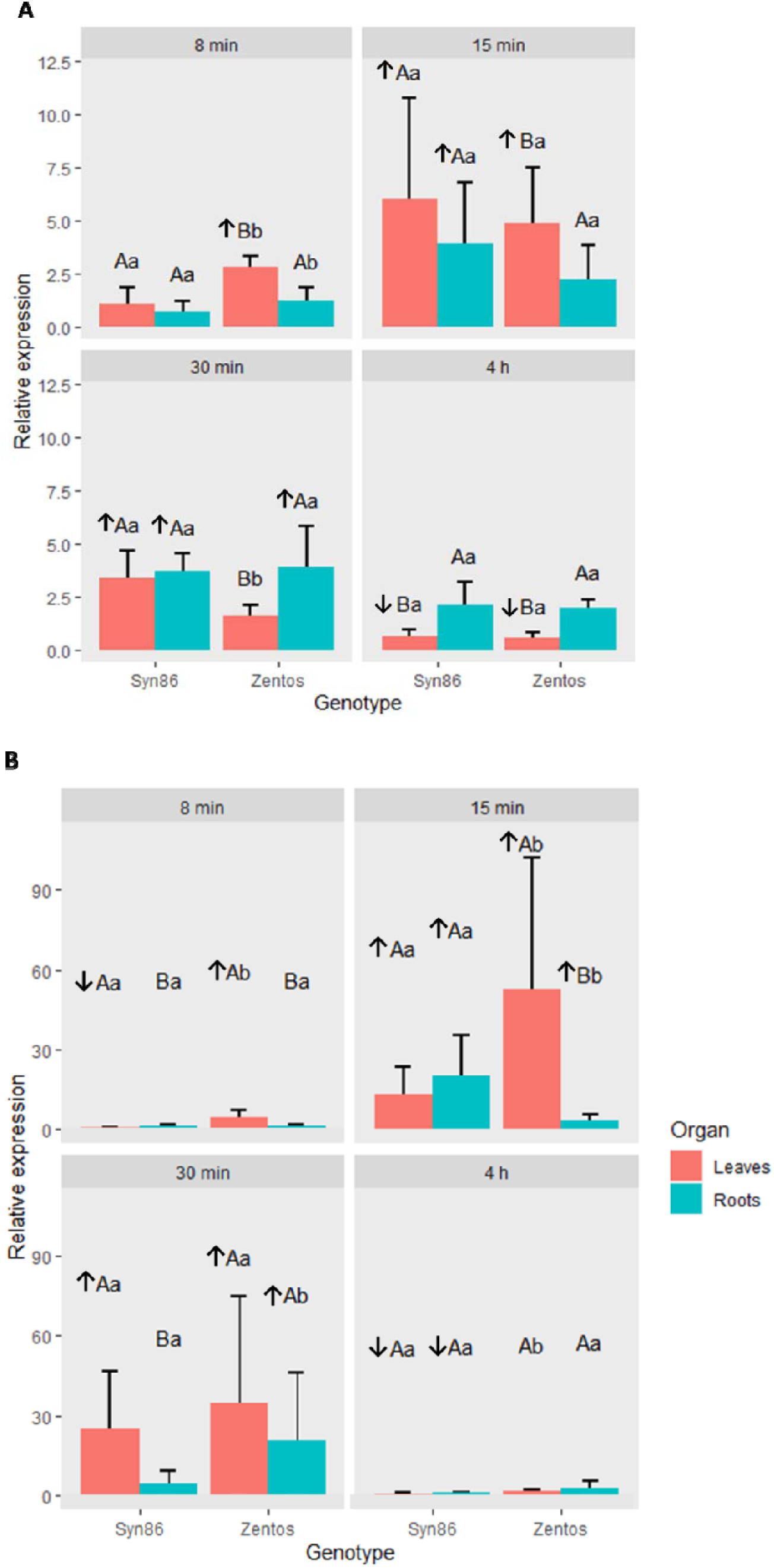
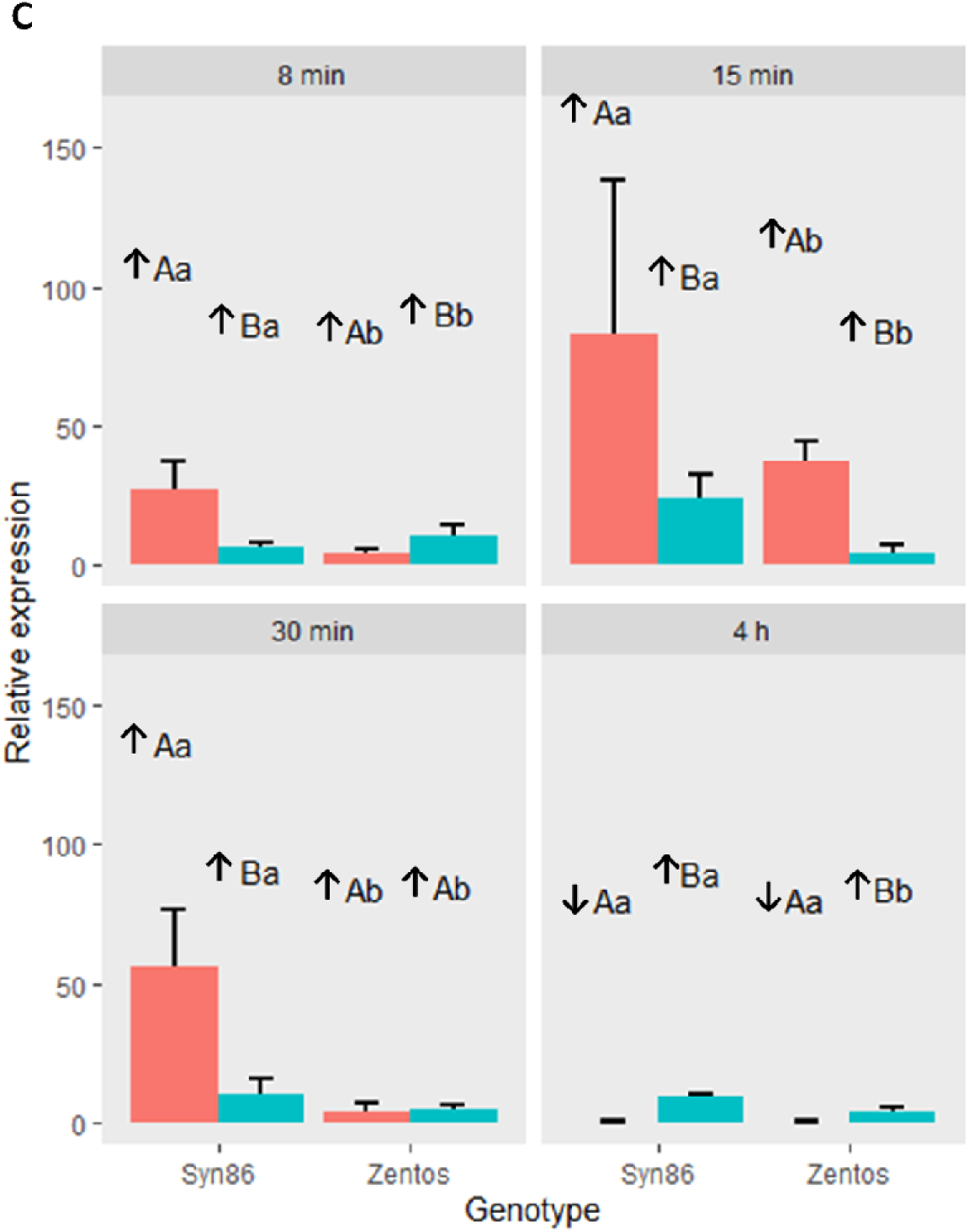
Relative expression values calculated with the ΔΔCt method (Livak and Schmittgen 2001) in leaves and roots from Syn86 and Zentos during the osmotic phase. (A) *TraesCS4D02G324800*, (B)*TraesCS5B02G299000* and (C)*TraesCS5D02G238700* expression. Different lower-case letters show significant differences in the mean values from the two genotypes in the same organ (p < 0.05), while the upper-case letters represent differences in the mean values from the two organs in the same genotype (*p* < 0.05). Mean relative expression values > 2.0 or < 1.0 (*p* < 0.05) indicated up-regulation (↑) or down-regulation (↓) of genes, respectively. Expression of leaves in (C) was extracted from Duarte-Delgado et al. (2020).

The relative expression values from the other member of the RBOH family, *TraesCS5B02G299000*, were higher (Fig. 6B). The analysis in leaves evidenced the opposite regulation of the expression of this transcript in the contrasting genotypes at 8 min ASE, with up-regulation in Zentos and down-regulation in Syn86. Later, at 15 and 30 min ASE the gene was up-regulated in both genotypes. At 15 min, the tolerant genotype registered a statistically significant higher expression than the susceptible (Fig. 6B). In roots, this transcript was up-regulated in both genotypes at 15 min and the relative expression value was significantly higher in the susceptible one. Later at 30 min, the gene was only up-regulated in Zentos (Fig. 6B). The expression analysis at 4h unveiled the down-regulation of the gene in both organs from the susceptible genotype, while in the tolerant it was not stress-responsive.

The study of *TraesCS5D02G238700* in roots complemented the analysis performed in leaves (Duarte-Delgado et al. 2020) and indicated the up-regulation of the transcript across all time points with mean relative expression values differing significantly in both genotypes. The mean expression values calculated for the organs were significantly different in all time points, except for Zentos at 30 min. At 15 min ASE, Syn86 registered the highest mean relative expression value (Fig. 6C).

### Diversity of salt-responsive transcription factors during the osmotic phase

To gain insights into the interplay of calcium-related signaling pathways and transcriptional regulation during stress response, the expression profiles from the salt-responsive TF families were characterized in Syn86 and Zentos. This analysis revealed the salt stress effect on the up-regulation of members from the AP2/ERF, WRKY, bZip, and GATA families in the two genotypes. The families HD-Zip, HSF, SFL, and MADS were down-regulated in Syn86 at 15 or 30 min ASE. Most TFs were up-regulated in Zentos at 15 min and presented greater relative expression values than those observed in Syn86 (Fig 7). AP2/ERF and WRKY were the most expressed families in the two genotypes (Fig 7). The expression profiles from the AP2/ERF family in Syn86 showed two peaks, with up-regulated genes at 8 and 30 min. On the other side, the members of this family in Zentos revealed higher expression values during one peak at 15 min (Fig 7). The members of the WRKY family presented an up-regulation peak at 30 min in both genotypes, with the greatest expression values in the tolerant. At 4 h ASE the down-regulation of these genes is observed in Syn86 (Fig 7). The members of the GATA family were up-regulated at 15 min in Zentos, and later at 30 min in Syn86. The fitted curves evidenced the up-regulation of members of the bZip family at 4 h ASE in both genotypes and Zentos unveiled far-reaching relative expression values. Finally, the HSF family revealed opposite expression values in the contrasting genotypes, with down- and up-regulation in Syn86 and Zentos, respectively (Fig 7). The relative expression values of the salt-responsive TF family members during the osmotic phase are listed in Supplementary Table S3.

**Fig. 7.**
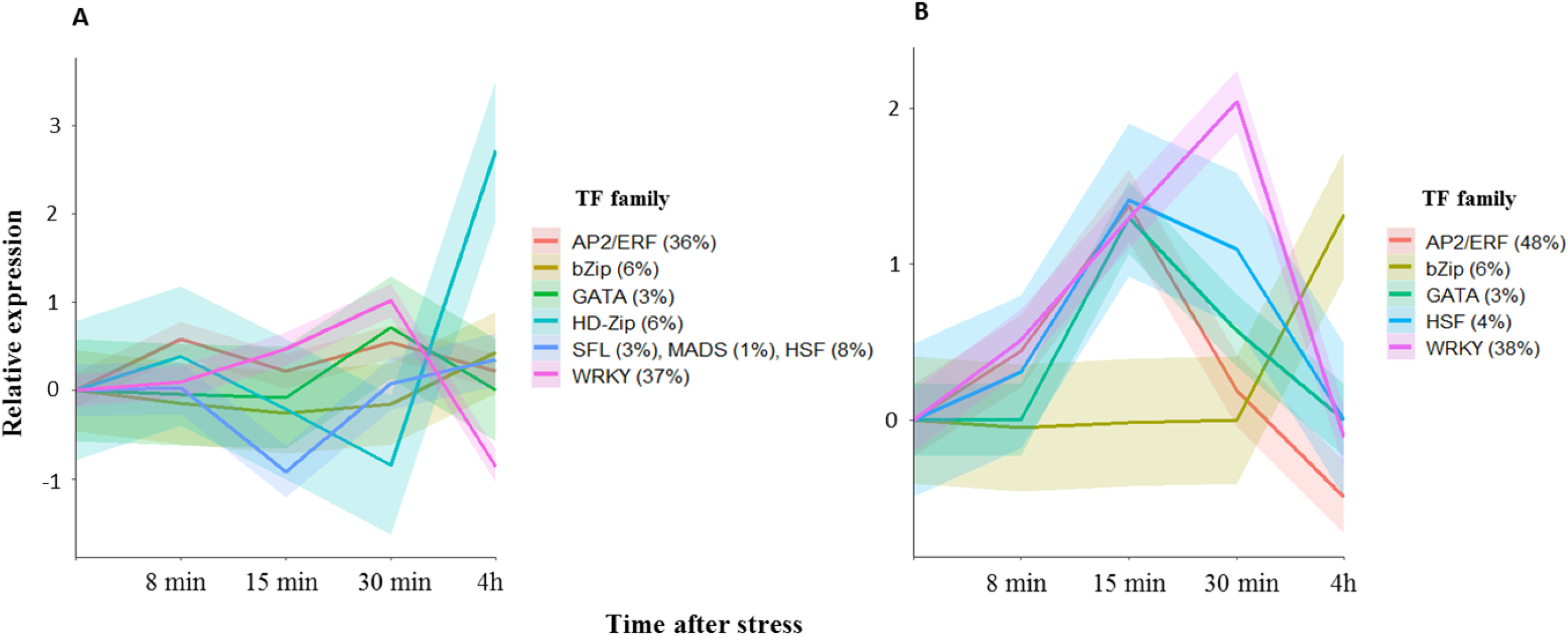
Expression patterns of salt-responsive transcription factor (TF) families in Syn86 (A) and Zentos (B) during the osmotic phase. The frequencies of the families are indicated in parentheses. A curve was fitted to represent the expression tendency of the members of each TF family and the shadows indicate the standard error of the relative expression values.

A comparative phylogenetic analysis of the salt-responsive WRKY and AP2/ERF genes facilitated their assignation to defined subfamilies to better understand their roles in the salt stress response. From 65 WRKY genes, 54% were expressed in both genotypes and 34% were exclusive from Syn86. Subfamily I contained 39% of the genes followed by subfamilies III (32%) and II (29%) (Supplementary Fig S2). Within the later subfamily, the IIc harbored most of TFs (23%), while only three genes were identified in group IId (5%).

The analysis of 84 salt-responsive AP2/ERF transcripts assigned 57% of them to the ERF (Ethylene-Responsive-Element-Binding protein) subfamily and 38% to the DREB (Dehydration Responsive Element-Binding) subfamily (Supplementary Fig S3). The ERF subfamily included 44% of transcripts with specific expression in Syn86 and 35% that were expressed in both genotypes. Among the salt-responsive DREB genes, 53% were expressed exclusively in Zentos followed by 28% of them expressed in both genotypes. *TraesCS2B02G542400*, *TraesCS5A02G473800*, *TraesCS1A02G058400* were assigned to the AP2 (APETALA2) subfamily according to the classification proposed by Zhao et al. (2019) for bread wheat. *TraesCS1B02G392300* was the only member from the RAV (Related to ABI3/VP) subfamily (Supplementary Fig S3).

### Identification of transcription factor binding sites

The analysis of the promoter regions of three selected calcium-binding genes was performed by amplicon sequencing in the two genotypes. More polymorphisms, 14 SNPs and two deletions of single nucleotides, were detected in *TraesCS5B02G299000* (Fig. 8A). A deletion of seven nucleotides in Syn86 and three SNPs were found in *TraesCS2D02G173600* promoter (Fig. 8B), while for *TraesCS5D02G238700* one single nucleotide insertion and two SNPs were identified in Zentos (Fig. 8C). The positions of the polymorphisms in the promoter sequences are shown in the Supplementary File S1. The TF binding site analysis predicted 17 families with potential binding abilities in regions adjacent to polymorphisms. From these families, bZip and C_2_H_2_ were common in the analyzed promoters (Fig. 8). The identification of bZip, GATA, ERF, and HD-Zip binding sites in the promoters (Fig. 8) is in line with the results obtained from the transcriptomics analysis where members from these families up-regulated under salt stress conditions were identified (Fig. 7).

**Fig. 8.**
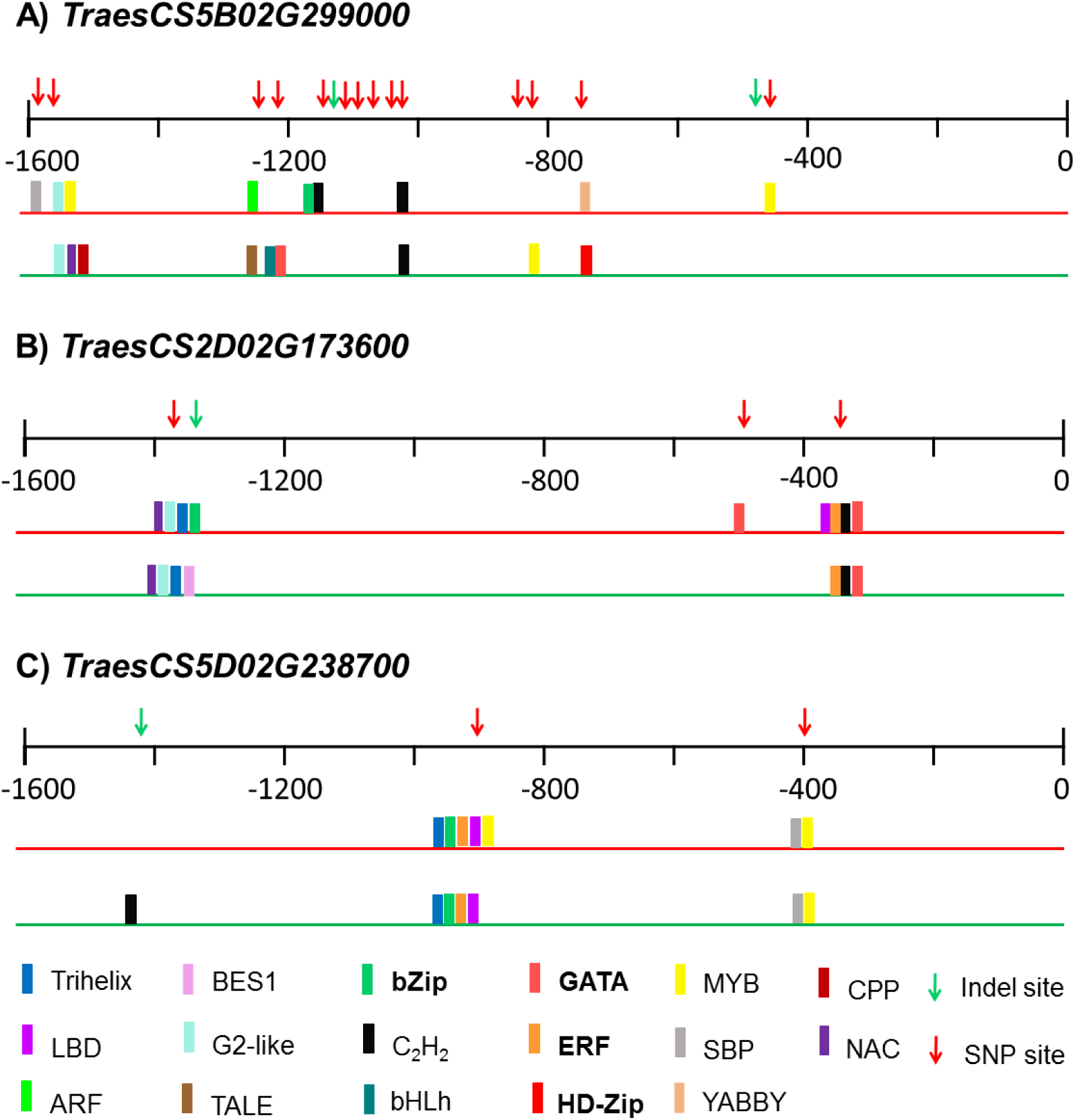
Transcription factor families with predicted binding capabilities to the polymorphic regions identified in the promoters of calcium-binding genes (A, B, and C) from Syn86 (red lines) and Zentos (green lines). Families with bold letters are salt stress-responsive as indicated in Fig. 6. The binding prediction was performed with the PlantTFDB (v 5.0) database (Jin *et al*., 2017).

### miRNA binding analysis based on SNPs identified in MACE reads

SNPs in calcium-binding genes were identified in the set of MACE reads previously collected from the contrasting genotypes (Duarte-Delgado et al. 2020), and miRNA binding sites were inferred in the polymorphic sites. A total of 82 SNPs were scored in 47 of the salt-responsive calcium-binding genes studied (Supplementary file S1). Most of them were located in 3’-UTR regions (52) followed by exons (21), six others were identified in novel 3’-UTR regions detected by Duarte-Delgado et al. (2020), and three in intron sequences. The surrounding sequence of eight SNPs in six genes contained potential miRNA binding sites (Table 3). From these genes, three were salt-responsive in Syn86, two in Zentos, and one in both genotypes. The possible effects of the SNP alternative allele in the miRNA affinity are shown in Table 3 and are described as the creation/loss of binding sites or increased/reduced complementarity. Five of the miRNAs found with interactions have been reported under biotic or abiotic stress stimuli in previous studies as presented in Table 3. For instance, responsive to salt and drought stresses in wheat is reported miR171a (Alptekin et al. 2017). This miRNA might bind to the gene sequence to regulate the expression of *TraesCS5B02G428400* in Zentos.

**Table 3.** Overview of the SNPs identified in salt-responsive calcium-binding genes with potential miRNA binding in the adjacent sequence to the polymorphisms predicted with the psRNAtarget tool (Dai et al. 2018).

| Gene | SNP<br>Position | Alternative<br>Allele<sup>a</sup> | Location<br>in gene | Salt-responsive<sup>b</sup> | miRNA binding<sup>c</sup> | Effect alternative<br>allele | Stimulus |
| --- | --- | --- | --- | --- | --- | --- | --- |
| *TraesCS1A02G239600* | 426881024 | T | 3'-UTR | Syn86 ↓ ↑ | ath-miR861-5p | loss of binding site | Nitrogen starvation (Liang et al. 2012) |
|  | 426881030 | A |  |  |  |  |  |
| *TraesCS1B02G251900* | 444688564 | T | 3'-UTR | Syn86 ↓ ↑ | ath-miR472-5p | loss of binding site | Insect and bacteria (Barah et al. 2013) |
|  |  |  |  |  | ath-miR851-3p | increased complementarity | Nitrogen starvation (Liang et al. 2012) |
| *TraesCS6B02G227900* | 354955972 | G | intron | Syn86 ↓ | ath-miR3933 | creation of binding site | Not reported |
| *TraesCS2A02G166200* | 118488650 | G | 3'-UTR | Zentos ↓ ↑ | ath-miR5649a | creation of binding site | Not reported |
|  |  |  |  |  | ath-miR5649b |  |  |
| *TraesCS5B02G428400* | 604067974 | G | 3'-UTR | Zentos ↑ | tae-miR5050 | reduced complementarity | Not reported |
|  | 604068120 | A | 3'-UTR |  | tae-miR171a | loss of binding site | Drought (Alptekin et al. 2017),<br>salt (Wang et al. 2014) |
|  |  |  |  |  | ath-miR171a-3p |  |  |
| *TraesCS7B02G350200* | 607529797 | G | exon | Both ↓ ↑ | ath-miR861-3p | reduced complementarity | Not reported |
|  |  |  |  |  | ath-miR775 | creation of binding site | Hypoxia (Moldovan et al. 2010) |

<sup>a</sup> Bold nucleotides were found in Zentos
<sup>b</sup> Up-regulation (↑) or down-regulation (↓) of genes
<sup>c</sup>ath: *A. thaliana*, tae: *T. aestivum*

## Discussion

### The calcium-binding-related transcriptional landscape in the contrasting genotypes revealed genes involved in rapid systemic ROS production

This study depicts the calcium-binding-related transcriptional landscape in bread wheat leaves and evidences a wide diversity of genes containing distinct types of architectures and domains in the contrasting genotypes. This diversity influenced the triggering of specific stress responses in systemic tissues during the osmotic phase (Gilroy et al. 2016). The domain analysis of the salt-responsive calcium-binding genes indicated that there are different types of them in the two genotypes. Zentos contained mainly genes corresponding to the EF-hand type, while Syn86 from the non-EF-hand one. The variation in the categories of stress-responsive genes involved in Ca^2+^- dependent networks on the contrasting genotypes represents a differential execution of signal transductions and suggests a particular calcium signature that can be related either to tolerance or susceptibility responses during the osmotic phase (Sanyal et al. 2019; Mohanta et al. 2019).

In addition to the transcripts of the OEC from the photosystem II related to photosynthesis reduction in Syn86 that were previously discussed (Duarte-Delgado et al., 2020), we highlighted additional genes within the non-EF-hand category associated with the susceptibility response from the genotype. For instance, annexins are plasma membrane proteins activated by ROS to function as Ca^2+^ channels (Moinoddini et al. 2023). The observed down-regulation of annexins could alter the transport of Ca^2+^ and affect the [Ca^2+^] cytosolic signals to be decoded by intracellular sensors (Liao et al. 2017; Sanyal et al. 2019). The up-regulation of CNX/CRT transcripts can be related to the greater oxidative stress response observed at the transcriptional level in this genotype (Duarte-Delgado et al. 2020). This calcium-binding protein type is involved in the pathway from the endoplasmic reticulum to refold and degrade stress-damaged proteins (Garg et al. 2015). We also identified the up-regulation of phosphoglycolate phosphatases, which corresponded to the EF-hand group. This transcriptional response from the susceptible genotype might be linked to the activation of the photorespiration metabolism to alleviate photoinhibition for stress protection (Wingler et al. 2000; Schwarte and Bauwe 2007). Thus, we could identify transcriptional responses involved in calcium signaling related to susceptibility responses.

Genes with EF-hand domains that detect transient increases of cytosolic Ca^2+^ and activate proper stress responses (Sanyal et al. 2019), were further studied through a phylogenetic analysis. Wheat CPKs and RBOHs showed clear orthologous relationships with the corresponding Arabidopsis proteins. The lack of clear orthologs for CaM and CML indicates greater sequence divergence along evolutionary time, and therefore gene function might be less conserved in distant taxa (Eisman and Kaufman 2013). The sequence similarity relatedness among the two species contributed to infer some stress response pathways in which the salt-responsive genes from Syn86 and Zentos might be involved.

For instance, the RBOH family members produce localized signaling ROS bursts and were of special interest in our study due to their involvement in rapid systemic stress responses (Choudhury et al. 2017; Chapman et al. 2019). The clustering with high statistical support of two wheat proteins with RBOHD from Arabidopsis is consistent with the reported role in salt stress response, systemic acquired acclimation by abiotic stress, and regulation of ABA-mediated stomatal closure (Chapman et al. 2019; Kaya et al. 2019). The RT-qPCR analysis in Zentos revealed the up-regulation of both RBOH genes in leaves at 8 min ASE, which coincided with a faster calcium and ROS signaling that is proposed as a crucial mechanism to trigger salt tolerance (Ismail et al. 2014). The higher expression levels of *TraesCS5B02G299000* in Zentos can be responsible to trigger tolerance mechanisms. Contrary to our study, Arabidopsis RBOHD was up-regulated exclusively in roots starting at 3 h ASE, and afterward RBOHE and RBOHF were expressed in leaves (Suzuki et al. 2011). These results suggest that a different RBOH member might regulate the onset of stress sensing in wheat roots at an earlier time point. The co-expression of RBOHD ortholog *TraesCS5B02G299000* and two CPKs from subgroup I in Zentos at 15 min, agrees with the kinase-substrate interactions from members from this subgroup and RBOHD in Arabidopsis to regulate TFs under stress (Yip-Delormel and Boudsocq 2019). Further studies at the protein level are necessary to validate these possible phosphorylation-mediated interactions in wheat to activate stress responses.

### Calcium-binding genes are predominantly co-expressed with WRKY and AP2/ERF members

This study suggests an important role of AP2/ERF and WRKY families on the osmotic phase response, as members from these families were the most abundant among the up-regulated TFs. According to studies in Arabidopsis, the simultaneous expression of the AP2/ERF family and EF-hand genes is intertwined. For instance, RBOHD expression is activated by the binding of ERF74 in the promoter region (Xie et al. 2019) and the transcription of some DREB genes from the AP2/ERF family is regulated by calmodulin-related pathways (Jan et al. 2017). Genes corresponding to the DREB subfamily showed greater relative expression values in the tolerant genotype, which can be related to the predominant activation of the ABA-independent signal transduction pathway during stress (Erpen et al. 2018; Li et al. 2019).

The fast expression of WRKY genes identified during the osmotic phase can contribute to the transcriptional activation of adaptation mechanisms related to ABA signaling and the production of secondary metabolites with relevance on stress responses (Banerjee and Roychoudhury 2015; Phukan et al. 2016). The co-expression of WRKY and GATA TFs identified in our study agrees with the transcripts identified in Arabidopsis after a few minutes of light stress activated by ROS/Ca^2+^ signals (Zandalinas et al. 2019). The three salt-responsive WRKY genes from Group IId detected in bread wheat might interact with EF-hand genes. In *A. thaliana* a CaM-binding domain has been identified and validated for this group of WRKY genes (Banerjee and Roychoudhury 2015; Seifikalhor et al. 2019). These observations support a fine-tuned interplay among calcium-binding genes and specific TF families in pathways to trigger salt stress responses.

### Genetic variation with potential influence on expression levels of calcium-binding genes

Several TF families showed potential binding sites adjacent to the identified polymorphisms in the promoter regions of the calcium-binding genes studied. Among them, the families GATA, bZip, HD-Zip, and ERF/AP2 were identified as salt-responsive in the comparative transcriptome analysis, highlighting them as putative regulators of differential expression of the calcium-binding genes in the contrasting genotypes. These expression regulations are supported by studies such as the one reporting the binding of a bZip member to the promoter of a calmodulin gene in Arabidopsis (Reddy et al. 2011). The identification of *cis*-regulatory polymorphisms in calcium-binding genes with affinity to ERF/AP2 and GATA families can underlie the co-expression of both gene categories that was discussed in the previous section. Interestingly, the polymorphisms detected in the promoter from *TraesCS2D02G173600*, a candidate gene identified within a salt stress response QTL (Dadshani 2018), can explain the up-regulation of this CML gene in Zentos (Duarte-Delgado et al. 2020) and the greater tolerance to salt stress found in this genotype.

The analysis of MACE reads led to the identification of SNPs to infer putative miRNA binding sites in the transcripts. Plants can respond to environmental stresses by altering gene expression through the activity of miRNAs (Khraiwesh et al. 2012; Asefpour Vakilian 2020). The potential interaction of the CPK transcript *TraesCS5B02G428400* and tae-miR171a agrees with the report of this miRNA as salt- and drought-responsive in wheat and other Triticeae species (Wang et al. 2014; Alptekin et al. 2017). The CPK transcript is stress-responsive for a short period, which can be related to tissue- and time-specific expression of miRNAs after salt stress exposure (Asefpour Vakilian 2020). Few SNPs were scored in intron regions and extended gene models described by Duarte-Delgado et al. (2020). The reads mapping to intronic regions can be due to imprecise annotations and they can be a proxy of alternative transcription and splicing events (Gaidatzis et al. 2015).

## Conclusions

The comprehensive analysis of the calcium-binding-related transcriptional landscape at the initial minutes and hours of the osmotic phase during the salt stress response revealed differences underlying the contrasting stress responses of the studied genotypes. The non-EF-hand category was specific for the susceptibility response, as revealed by the regulation of transcripts related to oxidative stress protection and oxidative stress damage repair. On the other side, Zentos, the tolerant genotype was characterized by a faster and higher up-regulation of EF-hand genes and TF members. The transcription of these genes might be involved in signaling pathways to trigger and increase salt tolerance, such as the early expression of RBOHD orthologs involved in ROS production. This study provides insights into the interplay of calcium-binding genes, WRKY, and AP2/ERF TF families in signaling pathways at the start of the osmotic phase to affect the expression of several downstream genes. Furthermore, the identification of natural variation in *cis-*regulatory sequences provides insights into mechanisms related to the differential expression levels observed among the contrasting genotypes, which can support the effect of QTL regions related to stress response.

## Acknowledgments

The authors want to acknowledge the staff from INRES Greenhouse Service for their support in the maintenance of the hydroponic experiments.

## Author contributions

DDD carried out the hydroponics and laboratory experiments, bioinformatics analyses, interpreted the data, and drafted the manuscript. SD guided and supported the hydroponics experiments. IV supported the hydroponics and RT-qPCR experiments, produced, and analyzed some amplicon sequences. AB and JL conceptualized the study and performed project administration. AB supervised the entire research, participated in the design of the experiments, and with the interpretation of the data. All authors participated in editing the manuscript and approved the final version.

## Data availability

The data supporting the results of this article are included within the article and the provided supplementary files.

## References

Ackermann M, Sikora-Wohlfeld W, Beyer A (2013) Impact of natural genetic variation on gene expression dynamics. PLoS Genet 9:e1003514.

Alaux M, Rogers J, Letellier T, Flores R, Alfama F, Pommier C, Mohellibi N, Durand S, Kimmel E, Michotey C, Guerche C, Loaec M, Lainé M, Steinbach D, Choulet F, Rimbert H, Leroy P, Guilhot N, Salse J, Feuillet C, Paux E, Eversole K, Adam-Blondon A-F, Quesneville H, International Wheat Genome Sequencing Consortium (2018) Linking the International Wheat Genome Sequencing Consortium bread wheat reference genome sequence to wheat genetic and phenomic data. Genome Biol 19:111.

Alptekin B, Langridge P, Budak H (2017) Abiotic stress miRNomes in the Triticeae. Funct Integr Genomics 17:145–170.

Asefpour Vakilian K (2020) Machine learning improves our knowledge about miRNA functions towards plant abiotic stresses. Sci Rep 10:3041.

Banerjee A, Roychoudhury A (2015) WRKY proteins: signaling and regulation of expression during abiotic stress responses. Sci World J 2015:807560.

Barah P, Winge P, Kusnierczyk A, Tran DH, Bones AM (2013) Molecular signatures in *Arabidopsis thaliana* in response to insect attack and bacterial infection. PLoS ONE 8:e58987.

Borrill P, Harrington SA, Uauy C (2019) Applying the latest advances in genomics and phenomics for trait discovery in polyploid wheat. Plant J 97:56–72.

Bürstenbinder K, Möller B, Plötner R, Stamm G, Hause G, Mitra D, Abel S (2017) The IQD family of calmodulin-binding proteins links calcium signaling to microtubules, membrane subdomains, and the nucleus. Plant Physiol 173:1692–1708.

Cai M, Chen LS, Liu J, Yang C (2020) IGREX for quantifying the impact of genetically regulated expression on phenotypes. NAR Genom Bioinform 2:lqaa010

Chapman JM, Muhlemann JK, Gayomba SR, Muday GK (2019) RBOH-dependent ROS synthesis and ROS scavenging by plant specialized metabolites to modulate plant development and stress responses. Chem Res Toxicol 32:370–396.

Choudhury FK, Rivero RM, Blumwald E, Mittler R (2017) Reactive oxygen species, abiotic stress, and stress combination. Plant J 90:856–867.

Curtis T, Halford NG (2014) Food security: the challenge of increasing wheat yield and the importance of not compromising food safety. Ann Appl Biol 164:354–372.

Dadshani S (2018) Genetic and physiological characterization of traits related to salinity tolerance in an advanced backcross population of wheat. PhD Thesis. University of Bonn. https://bonndoc.ulb.uni-bonn.de/xmlui/handle/20.500.11811/7341.

Dai X, Zhuang Z, Zhao PX (2018) psRNATarget: a plant small RNA target analysis server (2017 release). Nucleic Acids Res 46:W49–W54.

Duarte-Delgado D, Dadshani S, Schoof H, Oyiga BC, Schneider M, Mathew B, Léon J, Ballvora A (2020) Transcriptome profiling at osmotic and ionic phases of salt stress response in bread wheat uncovers trait-specific candidate genes. BMC Plant Biol 20:428.

Duarte-Delgado D (2020) Insights into the salt stress adaptation mechanisms of bread wheat genotypes using a systemic approach. PhD Thesis. University of Bonn. https://hdl.handle.net/20.500.11811/8599.

Eisman RC, Kaufman TC (2013) Probing the Boundaries of Orthology: The Unanticipated Rapid Evolution of *Drosophila* centrosomin. Genetics 194:903–926.

Erpen L, Devi HS, Grosser JW, Dutt M (2018) Potential use of the DREB/ERF, MYB, NAC and WRKY transcription factors to improve abiotic and biotic stress in transgenic plants. Plant Cell, Tissue Organ Cult 132:1–25.

Gaidatzis D, Burger L, Florescu M, Stadler MB (2015) Analysis of intronic and exonic reads in RNA-seq data characterizes transcriptional and post-transcriptional regulation. Nat Biotechnol 33:722–729.

Galon Y, Finkler A, Fromm H (2010) Calcium-regulated transcription in plants. Mol Plant 3:653–669.

Garg G, S Y, Ruchi, G Y (2015) Key roles of calreticulin and calnexin proteins in plant perception under stress conditions: A Review. Adv Life Sci 5:18–26

Gilroy S, Białasek M, Suzuki N, Górecka M, Devireddy AR, Karpiński S, Mittler R (2016) ROS, Calcium, and Electric Signals: Key mediators of rapid systemic signaling in plants. Plant Physiol 171:1606–1615.

Gupta B, Huang B (2014) Mechanism of salinity tolerance in plants: Physiological, biochemical, and molecular characterization. Int J Genomics 2014:701596.

Hall TA (1999) BioEdit: a user-friendly biological sequence alignment editor and analysis program for Windows 95/98/NT. Nucleic Acid Symp Ser 41:95–98.

Himanen SV, Sistonen L (2019) New insights into transcriptional reprogramming during cellular stress. J Cell Sci 132:jcs238402.

Ho HL (2015) Functional roles of plant protein kinases in signal transduction pathways during abiotic and biotic stress. J Biodivers Biopros Dev 2:147.

Ismail A, Takeda S, Nick P (2014) Life and death under salt stress: Same players, different timing? J Exp Bot 65:2963–2979.

IWGSC (2018) Shifting the limits in wheat research and breeding using a fully annotated reference genome. Science 361:eaar7191.

Jan A, Hadi F, Midrarullah, Ahmad A, Rahman K (2017) Role of CBF/DREB gene expression in abiotic stress tolerance. A Review. Int J Hort Agric 2:1–12

Jin J, Tian F, Yang D-C, Meng Y-Q, Kong L, Luo J, Gao G (2017) PlantTFDB 4.0: toward a central hub for transcription factors and regulatory interactions in plants. Nucleic Acids Res 45:D1040–D1045.

Julkowska MM, Testerink C (2015) Tuning plant signaling and growth to survive salt. Trends Plant Sci 20:586–594.

Katoh K, Rozewicki J, Yamada K (2019) MAFFT online service: multiple sequence alignment, interactive sequence choice and visualization. Brief Bioinformatics 20:1160–1166

Kaya H, Takeda S, Kobayashi MJ, Kimura S, Iizuka A, Imai A, Hishinuma H, Kawarazaki T, Mori K, Yamamoto Y, Murakami Y, Nakauchi A, Abe M, Kuchitsu K (2019) Comparative analysis of the reactive oxygen species-producing enzymatic activity of Arabidopsis NADPH oxidases. Plant J 98:291–300.

Keshtehgar, A., Rigi, K. and Vazirimehr, M. (2013) Effects of salt stress in crop plants. Int J Agric Crop Sci 5:2863–2867.

Khraiwesh B, Zhu J-K, Zhu J (2012) Role of miRNAs and siRNAs in biotic and abiotic stress responses of plants. Biochim Biophys Acta 1819:137–148.

Klein D, Ricordi C, Pastori RL (2004) Quantification of ribozyme target RNA using real-time PCR. In: Sioud M (ed) Ribozymes and siRNA protocols. Humana Press, Totowa, NJ, pp 49–56.

Konstantinov DK, Zubairova US, Ermakov AA, Doroshkov AV (2021) Comparative transcriptome profiling of a resistant vs susceptible bread wheat (*Triticum aestivum* L.) cultivar in response to water deficit and cold stress. PeerJ 9:e11428.

Kumar S, Stecher G, Li M, Knyaz C, Tamura K (2018) MEGA X: Molecular evolutionary genetics analysis across computing platforms. Mol Biol Evol 35:1547–1549

Kunert A, Naz AA, Dedeck O, Pillen K, Léon J (2007) AB-QTL analysis in winter wheat: I. Synthetic hexaploid wheat (*T. turgidum* ssp. *dicoccoides* x *T. tauschii*) as a source of favourable alleles for milling and baking quality traits. Theor Appl Genet 115:683– 695.

Lange W, Jochemsen G (1992) Use of the gene pools of *Triticum turgidum* ssp. *dicoccoides* and *Aegilops squarrosa* for the breeding of common wheat (*T. aestivum*), through chromosome-doubled hybrids. Euphytica 59:213–220.

La Verde V, Dominici P, Astegno A (2018) Towards understanding plant calcium signaling through calmodulin-like proteins: A biochemical and structural perspective. Int J Mol Sci 19:331

Li C, Zhang W, Yuan M, Jiang L, Sun B, Zhang D, Shao Y, Liu A, Liu X, Ma J (2019) Transcriptome analysis of osmotic-responsive genes in ABA-dependent and - independent pathways in wheat (*Triticum aestivum* L.) roots. PeerJ 7:e6519.

Li H, Handsaker B, Wysoker A, Fennell T, Ruan J, Homer N, Marth G, Abecasis G, Durbin R, 1000 Genome Project Data Processing Subgroup (2009) The Sequence Alignment/Map format and SAMtools. Bioinformatics 25:2078–2079.

Liang G, He H, Yu D (2012) Identification of nitrogen starvation-responsive microRNAs in *Arabidopsis thaliana*. PLoS ONE 7:e48951.

Liao C, Zheng Y, Guo Y (2017) MYB30 transcription factor regulates oxidative and heat stress responses through ANNEXIN-mediated cytosolic calcium signaling in *Arabidopsis*. New Phytol 216:163–177.

Livak KJ, Schmittgen TD (2001) Analysis of relative gene expression data using real-time quantitative PCR and the 2(-Delta Delta C(T)) method. Methods 25:402–408.

Loginova DB, Silkova OG (2018) The genome of bread wheat *Triticum aestivum* L.: Unique structural and functional properties. Russ J Genet 54:403–414.

Malabarba J, Meents AK, Reichelt M, Scholz SS, Peiter E, Rachowka J, Konopka-Postupolska D, Wilkins KA, Davies JM, Oelmüller R, Mithöfer A (2021) ANNEXIN1 mediates calcium-dependent systemic defense in *Arabidopsis* plants upon herbivory and wounding. New Phytol 231:243–254.

Medvedev SS (2018) Principles of Calcium Signal Generation and Transduction in Plant Cells. Russ J Plant Physiol 65:771–783.

Mitchell AL, Attwood TK, Babbitt PC, Blum M, Bork P, Bridge A, Brown SD, Chang H-Y, El-Gebali S, Fraser MI, Gough J, Haft DR, Huang H, Letunic I, Lopez R, Luciani A, Madeira F, Marchler-Bauer A, Mi H, Natale DA, Necci M, Nuka G, Orengo C, Pandurangan AP, Paysan-Lafosse T, Pesseat S, Potter SC, Qureshi MA, Rawlings ND, Redaschi N, Richardson LJ, Rivoire C, Salazar GA, Sangrador-Vegas A, Sigrist CJA, Sillitoe I, Sutton GG, Thanki N, Thomas PD, Tosatto SCE, Yong S-Y, Finn RD (2019) InterPro in 2019: improving coverage, classification and access to protein sequence annotations. Nucleic Acids Res 47:D351–D360.

Mohanta TK, Yadav D, Khan AL, Hashem A, Abd Allah EF, Al-Harrasi A (2019) Molecular players of EF-hand containing calcium signaling event in plants. Int J Mol Sci 20:1476.

Moinoddini F, Mirshamsi Kakhki A, Bagheri A, Jalilian A (2023) Genome-wide analysis of annexin gene family in *Schrenkiella parvula* and *Eutrema salsugineum* suggests their roles in salt stress response. PLoS One 18:e0280246.

Moldovan D, Spriggs A, Yang J, Pogson BJ, Dennis ES, Wilson IW (2010) Hypoxia-responsive microRNAs and trans-acting small interfering RNAs in *Arabidopsis*. J Exp Bot 61:165–177.

Montenegro JD, Golicz AA, Bayer PE, Hurgobin B, Lee H, Chan C-KK, Visendi P, Lai K, Doležel J, Batley J, Edwards D (2017) The pangenome of hexaploid bread wheat. Plant J 90:1007–1013.

Oyiga BC, Ogbonnaya FC, Sharma RC, Baum M, Léon J, Ballvora A (2019) Genetic and transcriptional variations in *NRAMP-2* and *OPAQUE1* genes are associated with salt stress response in wheat. Theor Appl Genet 132:323–346.

Oyiga BC, Sharma RC, Baum M, Ogbonnaya FC, Léon J, Ballvora A (2018) Allelic variations and differential expressions detected at quantitative trait loci for salt stress tolerance in wheat. Plant Cell Environ 41:919–935.

Parihar P, Singh S, Singh R, Singh VP, Prasad SM (2015) Effect of salinity stress on plants and its tolerance strategies: a review. Environ Sci Pollut Res Int 22:4056–4075.

Pessarakli M, Szabolcs I (2019) Soil salinity and sodicity as particular plant/crop stress factors. In: Pessarakli M (ed) Handbook of plant and crop stress, Fourth edition. CRC Press, Taylor & Francis Group, Boca Raton, FL, pp 3–21

Phukan UJ, Jeena GS, Shukla RK (2016) WRKY Transcription Factors: Molecular Regulation and Stress Responses in Plants. Front Plant Sci 7:760.

Ranty B, Aldon D, Cotelle V, Galaud J-P, Thuleau P, Mazars C (2016) Calcium sensors as key hubs in plant responses to biotic and abiotic stresses. Front Plant Sci 7:327.

Reddy ASN, Ali GS, Celesnik H, Day IS (2011) Coping with stresses: roles of calcium- and calcium/calmodulin-regulated gene expression. Plant Cell 23:2010–2032.

Roy SJ, Negrão S, Tester M (2014) Salt resistant crop plants. Curr Opin Biotechnol 26:115– 124.

Sanyal SK, Mahiwal S, Pandey GK (2019) Calcium Signaling: A Communication Network that Regulates Cellular Processes. In: Sopory S (ed) Sensory Biology of Plants. Springer, Singapore, pp 279–309.

Schneider M, Shrestha A, Ballvora A, Léon J (2022) High-throughput estimation of allele frequencies using combined pooled-population sequencing and haplotype-based data processing. Plant Methods 18:34.

Schwarte S, Bauwe H (2007) Identification of the photorespiratory 2-phosphoglycolate phosphatase, PGLP1, in *Arabidopsis*. Plant Physiol 144:1580–1586.

Seifikalhor M, Aliniaeifard S, Shomali A, Azad N, Hassani B, Lastochkina O, Li T (2019) Calcium-signaling and salt tolerance are diversely entwined in plants. Plant Signal Behav 14:1665455.

Shen J, Zhang J, Zhou M, Zhou H, Cui B, Gotor C, Romero LC, Fu L, Yang J, Foyer CH, Pan Q, Shen W, Xie Y (2020) Persulfidation-based modification of cysteine desulfhydrase and the NADPH oxidase RBOHD controls guard cell abscisic acid signaling. Plant Cell 32:1000–1017.

Shi S, Li S, Asim M, Mao J, Xu D, Ullah Z, Liu G, Wang Q, Liu H (2018) The Arabidopsis Calcium-Dependent Protein Kinases (CDPKs) and their roles in plant growth regulation and abiotic stress responses. Int J Mol Sci 19:1900

Shi X, Ling H-Q (2018) Current advances in genome sequencing of common wheat and its ancestral species. Crop J 6:15–21.

Sultana N, Islam S, Juhasz A, Yang R, She M, Alhabbar Z, Zhang J, Ma W (2020) Transcriptomic study for identification of major nitrogen stress responsive genes in Australian bread wheat cultivars. Front Genet 11:583785

Suzuki N, Miller G, Morales J, Shulaev V, Torres MA, Mittler R (2011) Respiratory burst oxidases: the engines of ROS signaling. Curr Opi Plant Biol 14:691–699.

Tuteja N (2007) Mechanisms of high salinity tolerance in plants. In: Häussinger D, Sies H (eds) Methods in Enzymology. Academic Press, pp 419–438

Varshney RK, Bohra A, Roorkiwal M, Barmukh R, Cowling WA, Chitikineni A, Lam H-M, Hickey LT, Croser JS, Bayer PE, Edwards D, Crossa J, Weckwerth W, Millar H, Kumar A, Bevan MW, Siddique KHM (2021) Fast-forward breeding for a food-secure world. Trends Genet 37:1124–1136.

Võsa U, Esko T, Kasela S, Annilo T (2015) Altered gene expression associated with microRNA binding site polymorphisms. PLoS ONE 10:e0141351.

Walkowiak S, Gao L, Monat C, Haberer G, Kassa MT, Brinton J, Ramirez-Gonzalez RH, Kolodziej MC, Delorean E, Thambugala D, Klymiuk V, Byrns B, Gundlach H, Bandi V, Siri JN, Nilsen K, Aquino C, Himmelbach A, Copetti D, Ban T, Venturini L, Bevan M, Clavijo B, Koo D-H, Ens J, Wiebe K, N’Diaye A, Fritz AK, Gutwin C, Fiebig A, Fosker C, Fu BX, Accinelli GG, Gardner KA, Fradgley N, Gutierrez-Gonzalez J, Halstead-Nussloch G, Hatakeyama M, Koh CS, Deek J, Costamagna AC, Fobert P, Heavens D, Kanamori H, Kawaura K, Kobayashi F, Krasileva K, Kuo T, McKenzie N, Murata K, Nabeka Y, Paape T, Padmarasu S, Percival-Alwyn L, Kagale S, Scholz U, Sese J, Juliana P, Singh R, Shimizu-Inatsugi R, Swarbreck D, Cockram J, Budak H, Tameshige T, Tanaka T, Tsuji H, Wright J, Wu J, Steuernagel B, Small I, Cloutier S, Keeble-Gagnère G, Muehlbauer G, Tibbets J, Nasuda S, Melonek J, Hucl PJ, Sharpe AG, Clark M, Legg E, Bharti A, Langridge P, Hall A, Uauy C, Mascher M, Krattinger SG, Handa H, Shimizu KK, Distelfeld A, Chalmers K, Keller B, Mayer KFX, Poland J, Stein N, McCartney CA, Spannagl M, Wicker T, Pozniak CJ (2020) Multiple wheat genomes reveal global variation in modern breeding. Nature 588:277– 283.

Wan S, Wang W, Zhou T, Zhang Y, Chen J, Xiao B, Yang Y, Yu Y (2018) Transcriptomic analysis reveals the molecular mechanisms of *Camellia sinensis* in response to salt stress. Plant Growth Regul 84:481–492.

Wang B, Sun Y-F, Song N, Wei J-P, Wang X-J, Feng H, Yin Z-Y, Kang Z-S (2014) MicroRNAs involving in cold, wounding and salt stresses in *Triticum aestivum* L. Plant Physiol Biochem 80:90–96.

Wang Y, Tiwari VK, Rawat N, Bikram S. Gill, Naxin Huo, You FM, Devin Coleman-Derr, Yong Q. Gu (2016) GSP: a web-based platform for designing genome-specific primers in polyploids. Bioinformatics 32:2382–2383

Wingler A, Lea PJ, Quick WP, Leegood RC (2000) Photorespiration: metabolic pathways and their role in stress protection. Philos Trans R Soc Lond B Biol Sci 355:1517–1529

Xie Z, Nolan TM, Jiang H, Yin Y (2019) AP2/ERF transcription factor regulatory networks in hormone and abiotic stress responses in arabidopsis. Front Plant Sci 10:228

Yip-Delormel T, Boudsocq M (2019) Properties and functions of calcium-dependent protein kinases and their relatives in *Arabidopsis thaliana*. New Phytol 224:585–604.

Zandalinas SI, Sengupta S, Burks D, Azad RK, Mittler R (2019) Identification and characterization of a core set of ROS wave-associated transcripts involved in the systemic acquired acclimation response of *Arabidopsis* to excess light. Plant J 98:126– 141.

Zhao Y, Ma R, Xu D, Bi H, Xia Z, Peng H (2019) Genome-wide identification and analysis of the AP2 transcription factor gene family in wheat (*Triticum aestivum* L.). Front Plant Sci 10:1286.

Zheng Y, Xu X, Li Z, Yang X, Zhang C, Li F, Jiang G (2009) Differential responses of grain yield and quality to salinity between contrasting winter wheat cultivars. Seed Sci Biotechnol 3:40–43.

Zhu T, Wang L, Rimbert H, Rodriguez JC, Deal KR, De Oliveira R, Choulet F, Keeble-Gagnère G, Tibbits J, Rogers J, Eversole K, Appels R, Gu YQ, Mascher M, Dvorak J, Luo M-C (2021) Optical maps refine the bread wheat *Triticum aestivum* cv. Chinese Spring genome assembly. Plant J 107:303–314.

